# TissuePlot: A Multi-Scale Interactive Visualization Tool for Spatial Data

**DOI:** 10.1101/2024.08.14.607906

**Authors:** Muhammed Khawatmi, Heba Sailem

## Abstract

Visualization of spatial datasets is essential for understanding biological systems that are composed of several interacting cell types. For example, gene expression data at the molecular level needs to be interpreted based on cell type, spatial context, tissue type, and interactions with the surrounding environment. Recent advances in spatial profiling technologies allow measurements of the level of thousands of proteins or genes at different spatial locations along with corresponding cellular composition. Representing such high dimensional data effectively to facilitate data interpretation is a major challenge. Existing methods such as spatially plotted pie charts obscure underlying tissue regions and necessitate switching between different views for comprehensive interpretations. Here, we present TissuePlot, a novel method for visualizing spatial data. TissuePlot tackles the key challenge of visualizing multi-scale phenotypic data at molecular, cellular and tissue level in the context of their spatial locations. To this end, TissuePlot employs a transparent hexagon tesselation approach that utilizes object borders to represent cell composition or gene-level data without obscuring the underlying cell image. Moreover, we implement a multi-view interactive approach, to allow interrogating spatial tissue data at multiple scales linking molecular information to tissue anatomy and motifs. We demonstrate TissuePlot utility using mouse brain data from the Bio+MedVis Redesign Challenge 2024. Our tool is accessible at https://sailem-group.github.io/TissuePlot/.

## 1 Introduction

Understanding the spatial patterns of cell types and their molecular profiles is crucial for distinguishing between healthy and diseased states, identifying better therapeutic targets, as well as gaining insights into the processes that drive tissue differentiation and organ formation in development. Traditional single-cell profiling techniques necessitate tissue dissociation, which results in the loss of spatial context. However, recent advancements have enabled the measurement of molecular states within cells while preserving their spatial locations, which is crucial for contextualizing molecular activities [1, 2, 3].

Spatial biology datasets pose significant challenges for data visualization due to their high dimensionality. Each dataset consists of microscopic tissue images along with measurements of the expression levels of thousands of genes or proteins at various spatial locations. This complexity necessitates advanced visualization techniques to accurately interpret and analyze the data. For most spatial profiling technologies, spatial location corresponds to 5-10 cells where cellular composition at a certain spatial location is based on cell type deconvolution techniques [3]. This multidimensionality necessitates visualization techniques that can simultaneously represent multiple phenotypes while overlaying this information on the underlying tissue image.

## 2 Contributions

Here we present the development of TissuePlot, a novel visualization tool for spatial data and demonstrate its utility based on data generated by the Visium platform as provided by the Bio+MedVis Redesign Challenge @IEEE Vis 2024 [4]. The data measure the expression of 31,053 genes in the mouse brain over an evenly spaced grid of 3499 spots, with each spot containing 5-10 cells. Beyond gene activity measurements, this data set enables the deconvolution of various cell types using a set of reference marker genes [3]. Moreover, clustering can be applied to group spots with similar gene expression or cellular proportions. Our work is aimed at developing a solution for several related tasks to facilitate the interpretation of spatial biology data (Figure 1). Our contributions are as follows:

**Figure 1:**
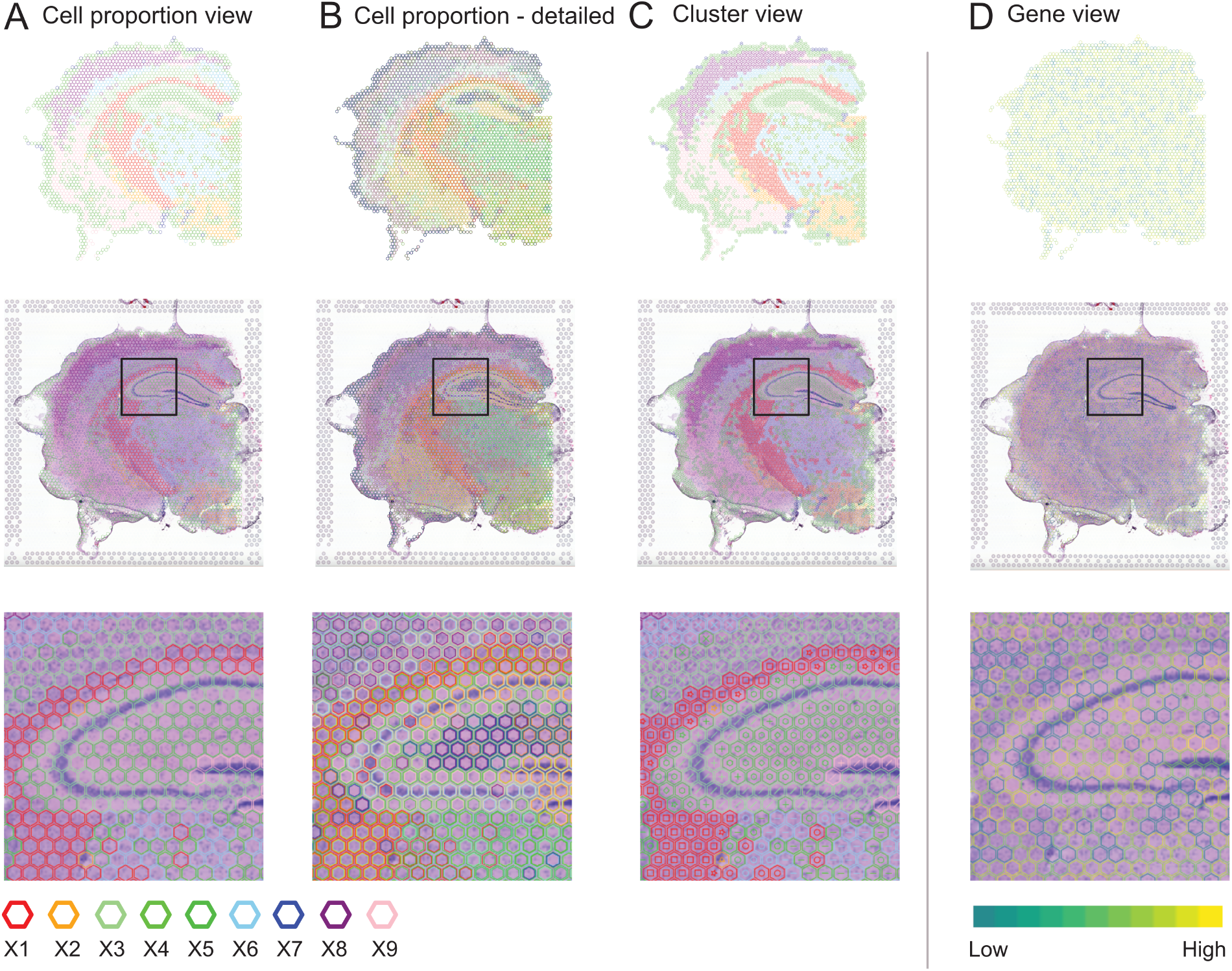
Different views in TissuePlot A-B) Cell proportions view with the most frequent cell type encoded to hexagon border color (A). B) Cell proportion detailed view where multiple outlined hexagons are layered to represent the top three most frequent cell types C) Cluster view where spot profiles are clustered and depicted as symbols. D) Gene view where gene levels are mapped to hexagon border color. TissuePlot representation with the tissue image displayed in the middle row and hidden in the top row. The bottom row provides a zoomed view of the area enclosed within the square in the middle row, highlighting a region of the hippocampus.

1. Introduce a novel method for representing spatial data using a transparent hexagon packing approach, where data is mapped to the color of the hexagon borders rather than their interiors. The primary advantage of this method is that it minimizes occlusion of the underlying tissue morphology. This approach can be applied to visualize cell composition through qualitative color mapping or gene expression levels through gradient-based color mapping, similar to a heatmap.
2. Propose interactive multi-scale visualization where additional layers of information are shown as the user zooms in. This enables the user to incrementally build different levels of understanding consistent with human learning [5]. At a high level, users can evaluate the most predominant cellular composition then zoom in to grasp other key cell types progressively.
3. Employ additional visual channels beyond color and position to add details on demand and create an implicit hierarchy. Specifically, users can also choose to plot the spot clusters as symbols at the centre of the hexagon, defined either based on cell composition profiles or gene expression profiles.
4. Develop an accessible web-app that allows users to visualize their own data (https://sailem-group.github.io/TissuePlot/). As all visualizations are performed on the client-side, data privacy and confidentiality concerns are eliminated.

## 3 Related Work

Existing methods for visualizing spatial data adapt traditional visualization methods such as heatmaps and pie charts to represent gene level information and tissue composition respectively. For example, a common approach is overlaying tissue regions with pie charts (Figure 2). However, these approaches do not allow the assessment of both tissue image and proportions of various cell types simultaneously, making it difficult to understand the complex spatial relationships within the tissue. Other approaches focused on dimensionality reduction to generate 2D scatter plots using tools such as those based on UMAP. For example, TissUUMAP, a desktop application generates a 2D latent space using UMAP and links this information to spatial locations via an interactive analysis interface [6]. These approaches focus is determining key phenotypes and overall structure of the data rather than the spatial tissue structure. Visualization of other aspects of spatial data, such as cell interactions, has been reviewed by Liu et al [2].

**Figure 2:**
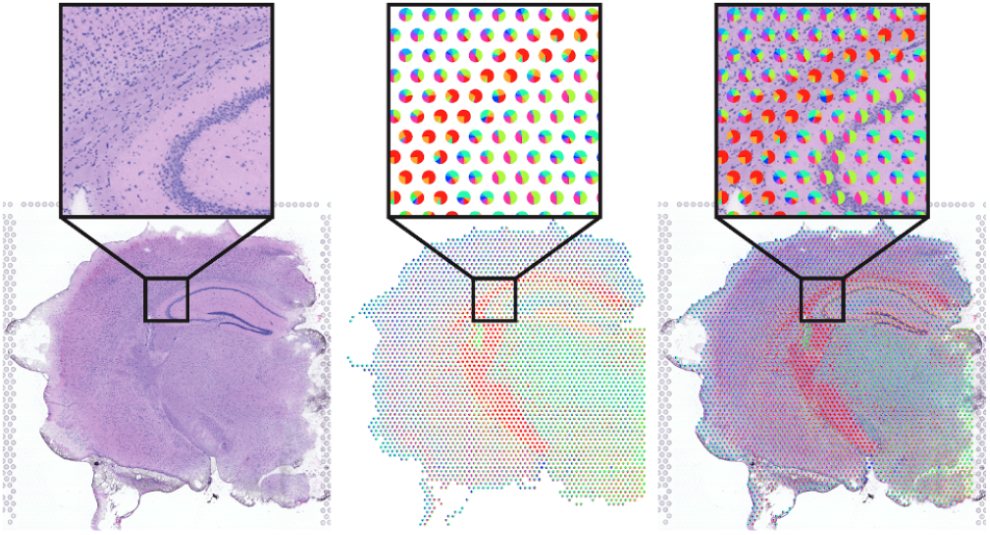
Pie chart representation of cell proportions provided by the Bio+MedVis Challenge 2024 [4].

Other glyph-based approaches for image data, such as Pheno-Plot and ShapoGraphy [7] allows intuitive visualization of multiple dimensions but are most effective for providing quantitative and pictorial summaries of the data. For example, to summarize key phenotypes identified in cell image and multiplexed data. However, they can be less effective at plotting large numbers of points in spatial datasets.

The use of hexagon maps for representing geographical and spatial data is well-established. Unlike circles, hexagons can tessellate seamlessly without leaving gaps, making them ideal for consistent spatial mapping [8]. Additionally, hexagons possess a natural visual appeal, often found in nature, such as in honeycomb structures ([9]). This shape reduces edge effects compared to square or triangle tessellations, offering a more effective method for aggregating spatial data. This not only enhances the visual accuracy of the representation but also improves the interpretability of the data, making hexagons a preferred choice in various geospatial and data visualization applications.

## 4 Dataset

We use data provided by Bio+MedVis Redesign Challenge 2024. Briefly, the data measure expression values of 31,053 genes in a section from a mouse brain for 3499 spots (i.e. spatial location) using Visium platform. A microscopic image of the same tissue section stained with Hematoxylin and Eosin is also captured.

Deconvolution methods were used to determine the proportion of different cell types present at each spot as each spot corresponds to mRNA from 5-10 cells [1]. These were determined based on gene expression profiles of known reference cell types based on single cell RNA-sequencing data. Through this analysis, nine key cell types, labeled X1-X9, were identified. Several spots were predominantly occupied by a single cell type, suggesting a more homogeneous tissue region (e.g., cell types X1 and X5 in Figure 3). Notably, most spots did not exhibit more than three cell types with a proportion exceeding 20%. This suggests that focusing on the three most common cell types could be a useful strategy.

**Figure 3:**
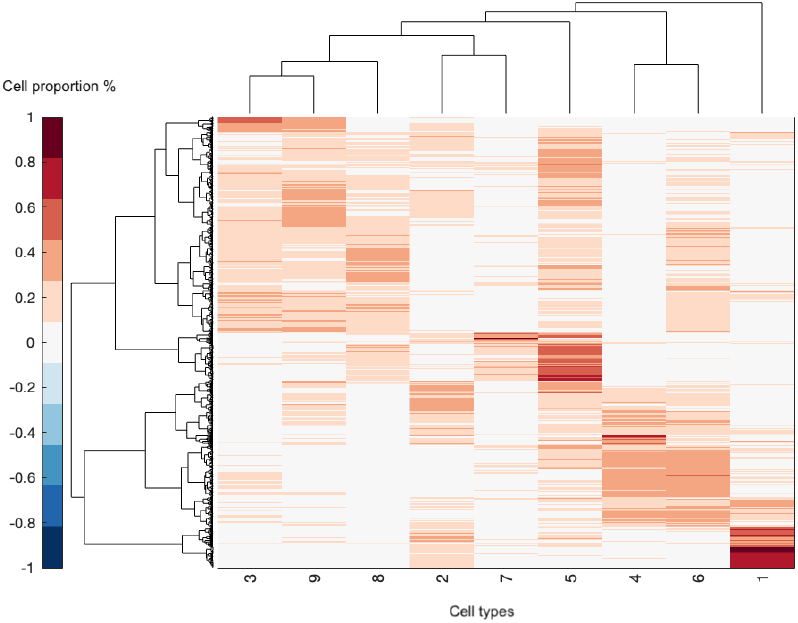
Visualization and clustering of cell proportion profiles for the 3499 spots. X-axis indicate cell types X1-X9.

Many genes were not detected in the sample or their expression did not vary a lot across spots. We identified the most expressed genes based on a standard deviation greater than 1.5. This resulted in 2138 genes available to view in our demo.

## 5 TissuePlot overview

### 5.1 Design principles

TissuePlot is an accessible web-based tool for visualizing spatial data. Its user-friendly interface is designed to be used by broad audience, including biologists and scientists with minimal computational experience.

TissuePlot overlays spatial measurements on top of the corresponding tissue images. The backbone of our representation employs outlined transparent hexagons that allow the underlying tissue image data to remain visible while mapping data to the hexagon borders.

We identify multi-scale visualization as a critical principle for effectively representing spatial data. This involves integrating tissue images, molecular information, and key phenotypes identified by clustering. To support this, we offer three views that enable users to progressively assess higher-level phenotypes and delve deeper into different tissue regions. These views include Cell proportions view, Cluster view, and the Gene view

Interactivity is another essential principle in our design. This feature is particularly useful during the early stages of data exploration, identification of key trends, and interpretation of analysis results from clustering or cell deconvolution approaches.

### 5.2 Visual channels

Our work employs three primary visual channels: color, position, and symbols. Each of these channels plays a crucial role in effectively visualizing large-scale biological data.

Color is a widely used channel for representing both categorical and continuous data. It scales efficiently with large datasets, making it ideal for distinguishing between different cell types (categorical data) and depicting varying levels of gene expression (continuous data). The use of color allows for immediate visual differentiation and interpretation of complex datasets. Combining color with other visual channels, such as utilizing tooltips, can help mitigate the limitations of the contextual nature of color.

Position. Spatial data demands utilization of this channel. By accurately mapping data points to their spatial locations, we can maintain the contextual relevance of the biological information, providing insights into the spatial relationships and interactions within the tissue samples.

Symbols are used to represent additional categorical variables, such as spot clusters. By assigning unique symbols to different clusters, we add another layer of information that can be easily interpreted alongside the color-coded data.

Importantly, the integration of these visual channels within their spatial context, combined with the use of multiple hexagons, establishes an implicit hierarchy that is more than the sum of these parts. This hierarchical structure not only enhances data interpretation by allowing users to discern high-level patterns but also facilitates detailed exploration of specific regions.

### 5.3 TissuePlot views

The following views allow overlaying various information (Figure 1), sometimes simultaneously as the user build mental maps of the data.

#### 5.3.1 Cell proportions view

Each spot is represented as a transparent hexagon where each cell type is assigned a distinct color. The color of the most predominant cell type at a certain location is mapped to the hexagon border color (Figure 1A).

Other frequently occurring cell types are displayed as inset hexagons, either on zooming in or on demand (Figure 1B). This creates a visual hierarchy where the outermost hexagon represents the most abundant cell type and the innermost hexagon represents the cell type with the lowest percentage among the three. We chose to display the top three cell types because each spot tends to be dominated by no more than three cell types (Section 4). An information box provides a detailed examination of the exact proportions of different cell types when the user hovers over a specific spatial location (Figure 4).

**Figure 4:**
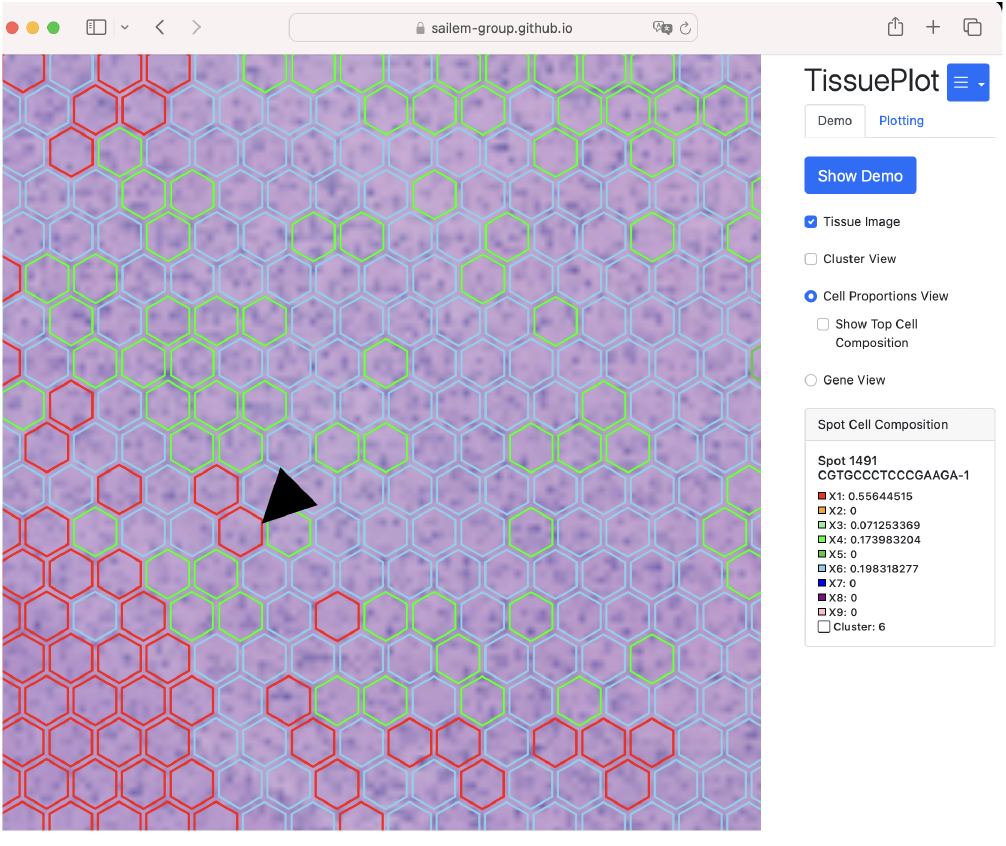
TissuePlot web interface with various views selectable from the right menu. The black arrow indicates the hovered hexagon, with the information box (bottom right) displaying the exact percentages of each cell type.

#### 5.3.2 Cluster view

We offer an additional view to represent spot clusters. Each cluster is represented by a distinct symbol, that is plotted at the center of each hexagon (Figure 1C). The color of each symbol is matched to the color of its corresponding hexagon for visual consistency and aesthetic appeal.

The clusters can be defined based on clustering of cell proportions, clustering of gene profiles, or any other grouping the user uploads. In our demo, cell proportion profiles were grouped into 10 clusters using k-means and 10 iterations. Importantly, this view can be combined with cell proportion view or gene expression view which further facilitates the interpretation of spot information.

#### 5.3.3 Gene expression view

Similar to cell proportion representation, but hexagon border colors are mapped to the log-transformed gene values based on a gradient color map. In this view, the user can view the expression values of one gene at a time. For our demo, we selected the top most expressed genes (Figure 1D).

### 5.4 Implementation

TissuePlot is developed using javascript and p5.js library. Its client-side architecture ensures that all data processing and visualization occur directly within the user’s browser, safeguarding data privacy. Users can either upload their own data for analysis or explore interactive demo to understand the tool capabilities.

TissuePlot requires data to be processed in appropriate software, such as R and Python, before being uploaded to the web application including cluster analysis. This ensures efficient, lightweight operations within the website.

## 6 Visualization of mouse brain data using Tissue-Plot

First, we find that using hexagon border colors for mapping data achieves the same function as a filled hexagon without obscuring underlying image data. For instance, users can see the underlying anatomy of the hippocampus region in the brain while examining cell proportion or gene levels (Figure 1C). In the Cell proportion view, displaying only the most dominant cell type is an effective approach to allow users to identify the key tissue regions. Users can zoom-in to various regions to determine additional cell compositions. The information box provides exact percentages (Figure 4). Alternatively, the user can also show the most common cell types for a given spot in the zoomed-out view. However, this may make interpretation more challenging for some users due to the interaction of multiple colors. Our detailed information box can help mitigate this issue (Figure 4).

In addition to color, plotting spot clusters as symbols at the center of the hexagon, can facilitate tissue-level phenotypes. For example, the hippocampus region shown in Figure 1A-B exhibits variation in most dominant cell types. However, by using k-means clustering to group spots with similar cell proportion profiles, we can discern that the spots in this region could vary in their composition, as indicated by the ‘x’ and ‘/’ symbols in the top of the zoomed region (Figure 1C, bottom row). This representation requires only one mental operation, unlike comparing multiple sections in a pie chart (Figure 2). Therefore clustering provides a higher level of information that enables much more effective comparison between spots. Moreover, this approach improves our understanding of how cell composition diverges across different anatomical regions.

## 7 Discussion

Effective data visualization in biology is crucial for interpreting complex datasets and deriving meaningful insights. Interactive and flexible visualization tools that can handle the complexity and density of spatial data without overwhelming the user are essential for advancing our understanding of biological systems. These tools have been applied successfully to various types of biological data [10, 11].

TissuePlot combines various conceptual solutions to tackle some of these challenges. First, it minimizes occlusion of underlying tissue regions, associated with pie-chart representations, by encoding cell composition data into the hexagon border colors. Although this approach only allows the visualization of the most dominant cell types, it offers advantages by being less overwhelming and enabling users to discern larger patterns and structures in the tissue. This concept of providing an overview view and details on demand has been investigated in depth in the literature [12] and is well adopted in the genomics field [13, 14]. We believe our approach will assist in identifying patterns and relationships from complex data by viewing multiple structures simultaneously or providing contextual information upon zooming.

The second challenge TissuePlot addresses is the difficulty in comparing thousands of pie charts. We tackle this by clustering spots with similar cell proportion profiles. These clusters can be represented as symbols, providing finer details and sparing users from performing such comparisons mentally [5].

We anticipate that such functionalities would enhance the ability to interpret complex biological data, aiding researchers in gaining deeper insights into molecular mechanisms within tissues. Evaluation of such encodings and their applicability to other types of spatial data, such as those subcellular level, or geographical data are interesting future directions.

## Acknowledgments

We acknowledge all members of the Sailem group. HS is funded by a Wellcome Career Development Award 225974/Z/22/Z.

